# High developmental temperature leads to low reproduction despite adult temperature

**DOI:** 10.1101/2020.06.08.140277

**Authors:** Marta A. Santos, Ana Carromeu-Santos, Ana S. Quina, Mauro Santos, Margarida Matos, Pedro Simões

## Abstract

Phenotypic plasticity can be an important tool in helping organisms to cope with changing thermal conditions and it may show an interdependency between life-stages. For instance, exposure to stressful temperatures during development can trigger a positive plastic response in adults. In this study, we analyse the thermal plastic response of laboratory populations of *Drosophila subobscura*, derived from two contrasting latitudes of the European cline. We measured fecundity characters in the experimental populations after exposure to five thermal treatments, with different combinations of developmental and adult temperatures (14°C, 18°C or 26°C). We ask whether (1) adult performance is enhanced (or reduced) by exposing flies to higher (or lower) temperatures during development only; (2) flies raised at lower temperatures outperform those developed at higher ones, supporting the “colder is better” hypothesis; (3) there is a cumulative effect on adult performance of exposing both juveniles and adults to higher (or lower) temperatures; (4) there is any evidence for historical effects on adult performance. Our main findings show that (1) higher developmental temperatures led to low reproductive performance regardless of adult temperature, while at lower temperatures reduced performance only occurred when cold conditions were persistent across juvenile and adult stage; (2) flies raised at lower temperatures did not always outperform those developed at other temperatures; (3) there was no (negative) cumulative effect of exposing both juveniles and adults to higher temperatures; (4) both latitudinal populations showed similar thermal plasticity patterns. The negative effect of high developmental temperature on reproductive performance, regardless of adult temperature, highlights the developmental stage as a critical and most vulnerable stage to climate change and associated heat waves.

## 1. Introduction

Phenotypic plasticity can help organisms to cope with climate change, by allowing for a rapid response to changing thermal conditions. In particular, developmental thermal plasticity can confer the potential to mount beneficial responses through developmental acclimation – *i.e*. higher resistance in adults as a consequence of exposure to stressful temperatures during development (Beaman et al., 2016; Sgrò et al., 2016; Sørensen et al., 2016). Plastic responses in insects have received much attention in recent years, particularly those associated with physiological tolerance to either cold or heat extremes. Studies in this context reveal a high potential for beneficial cold acclimation while heat acclimation is likely physiologically constrained and potentially subjected to trade-offs (e.g. (Kellermann et al., 2012; Kellermann and Sgrò, 2018; MacLean et al., 2019; Schou et al., 2017).

An increasing number of experiments is starting to address more thoroughly the developmental thermal plasticity associated with adult life-history traits, mainly in ectotherms (Angilletta et al., 2019; Austin and Moehring, 2019; Cao et al., 2018; Klepsatel et al., 2019; Klockmann et al., 2017; Kristensen et al., 2012; Manenti et al., 2017; Porcelli et al., 2017; Zamorano et al., 2017). This is extremely relevant as traits closely related to fitness, such as fecundity and longevity, are crucial for population persistence and likely to be affected by climate change (Walsh et al., 2019). In general, evidence was found for a negative impact of high developmental temperature on adult fitness (e.g. (Cao et al., 2018; Klepsatel et al., 2019; Klockmann et al., 2017; Porcelli et al., 2017). On the one hand, some studies addressing fecundity patterns support the “optimal developmental temperature” hypothesis, where individuals developed at intermediate, “optimal” temperatures show general better adult performance across environments relative to individuals developed at more extreme temperatures (Klepsatel et al., 2019; Kristensen et al., 2012). On the other hand, evidence for the impact of colder developmental temperatures in fecundity is less conclusive, with positive effects observed in some studies (e.g. (Nunney and Cheung, 1997; Simões et al., 2020; Zamorano et al., 2017), but not in others (Angilletta et al., 2019; Huey et al., 1995; Klepsatel et al., 2019; Kristensen et al., 2012).

*Drosophila subobscura* is an excellent model organism to study thermal responses. This species shows latitudinal clinal variation for chromosomal inversion frequencies in three distinct continents, and thermal adaptation is likely the main driving force (Prevosti et al., 1988; Rezende et al., 2010). These polymorphisms have been shifting worldwide as a consequence of global warming (Balanyá et al., 2006) while also clearly responding to sudden heat waves (Rodríguez-Trelles et al., 2013). Adult thermal plasticity with a beneficial response for thermal tolerance under colder, but not warmer, environments has been described in *D. subobscura* (MacLean et al., 2019; Schou et al., 2017). Developmental thermal plasticity has also been studied in this species with lower adult performance associated with higher developmental temperatures (Porcelli et al., 2017; Simões et al., 2020, see below). In addition, Porcelli et al. (2017) found evidence for geographical differences in the response to such thermal scenarios.

Our team has shown that populations founded from some extreme locations of the European cline (Portugal and The Netherlands) not only had a clear genetic response to new environmental conditions (Fragata et al., 2014; Simões et al., 2017), but also exhibited plastic responses to new thermal challenges even after 30+ generations of laboratory confinement (Fragata et al., 2016). We further addressed the developmental thermal plasticity in two sets of these historically differentiated populations after 67 generations of evolution at 18°C by subjecting juvenile and adult flies to different temperature combinations of 15 °C, 18°C, and 25°C (Simões et al., 2020). First, we observed that increased temperatures (25°C) during both juvenile and adult stages led to a very poor reproductive performance, while higher temperatures only in the adult stage increased adult reproductive performance. Second, we found that flies that developed at colder temperature (15°C) had increased reproductive performance at 15°C, when compared to those developed at 18°C (control conditions), indicating cold acclimation. Finally, we found some historical effect in thermal plasticity for fecundity, with a higher cold acclimation of the southern populations (Simões et al., 2020). The results suggested that heat has a negative effect during fly’s development. However, because the effect of developmental temperature was not assessed separately, we could not determine whether the low adult performance resulted from the combined negative effect of high developmental and adult temperatures, or the negative effect of high developmental temperature alone. Also, the cold acclimation pattern that we found can be the result of a general better performance of individuals raised at lower temperatures, perhaps because of a higher ovariole number (Moreteau et al., 1997). This “colder is better” hypothesis (Huey et al., 1999) can be tested by comparing adult performance of flies raised at low (15°C) and control (18°C) temperatures in a control environment.

To answer the issues raised by the Simões et al. (2020) study, we here report an additional developmental plasticity experiment in the same populations, where we test the adult performance of individuals exposed to three different temperatures (14°C, 18°C, and 26°C). Our main questions are:

1. Is adult performance enhanced (or reduced) by exposing flies to higher (or lower) temperatures during development only? And, more specifically, do flies raised at lower temperatures outperform those developed at higher ones – “colder is better” hypothesis (Huey et al., 1999)?
2. Is there a cumulative effect on adult performance of exposing both juveniles and adults to higher (or lower) temperatures?
3. Is there any evidence for historical effects on adult performance, particularly at lower temperatures (as previously found)?

## 2. Material and Methods

### 2.1 Origin and maintenance of Laboratory Populations

This study involved two sets of laboratory populations derived from independent collections in natural populations of *Drosophila subobscura* sampled in late August/early September 2013. These were done in two contrasting European latitudes: Adraga (Portugal) and Groningen (Netherlands) – see details in (Simões et al., 2020, 2017). Each latitudinal population was three-fold replicated after founding, giving rise to PT_1-3_ and NL_1-3_ populations. Population maintenance involved discrete generations with a synchronous 28-day cycle; 12L:12D photoperiod; constant temperature of 18°C; controlled densities in adults (50 adults per vial) and eggs (70 eggs per vial); reproduction for the next generation was around peak fecundity (seven to ten days old imagoes). Census size of populations ranged between 500 and 1200 individuals (see also (Simões et al., 2017). The thermal plasticity assay was performed when PT and NL populations were at their 71^st^ generation of evolution in the laboratory environment.

### 2.2 Thermal Plasticity assay

To study the effect of different thermal environments on adult performance, we analysed fecundity and wing size in PT and NL populations subjected to five thermal treatments (see Figure 1). In three of them, we exposed the flies to the same developmental and adult temperatures, 14°C, 18°C or 26°C (treatments 14-14, 18-18, and 26-26, respectively). The other two treatments were applied to analyse the impact of (higher or lower) developmental temperature on adult reproductive performance tested in control conditions. This was done by assaying imagoes at 18°C following their development at either 14°C or 26 °C (treatments 14-18 and 26-18, respectively). Twenty recently emerged mating pairs (virgin males and females) per population and treatment were formed, with a total of 600 pairs (20 pairs*6 populations*5 temperature treatments). Flies were transferred to fresh medium every other day, vials were daily checked for the presence of eggs, and the eggs laid by each female were counted between days 7 and 8 since emergence. This procedure allowed to estimate two adult parameters: age of first reproduction (number of days since emergence until the first egg laying) and fecundity (measured as the total number of eggs laid between days 7 and 8). The first parameter addresses the rate of sexual maturity, while the second refers to a period that is close to the age of egg collection for the following generation, where selective pressures are likely higher in all experimental populations.

**Figure 1.**
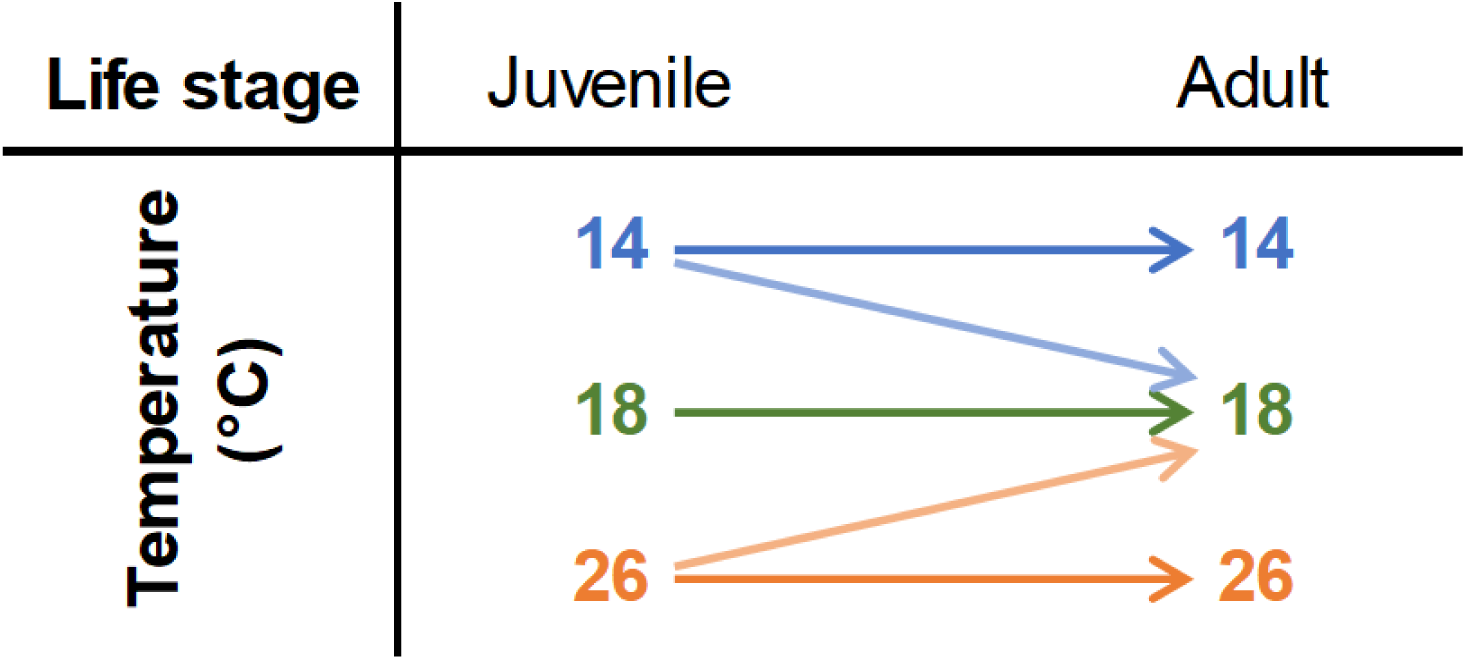
Experimental design applied, with three developmental and three adulthood temperatures.

### 2.3 Statistical Methods

Thermal plasticity was analysed by linear mixed models fitted with REML (restricted maximum likelihood). P-values for differences between temperatures, populations (PT or NL), and their interaction were obtained through analyses of variance (Type III Wald F tests, Kenward-Roger degrees of freedom). The model applied was as follows:

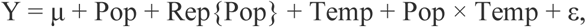

Y is the trait in study (age of first reproduction or fecundity), Pop is the fixed factor “latitudinal population” (with categories PT and NL), Rep{Pop} is the random factor replicate population nested in the fixed factor latitudinal population, and Temp is the fixed factor corresponding to the different temperature treatments. Raw data is the mean value for each replicate population and temperature treatment, *e.g*. NL_1_ for the 14-14 treatment. Higher fecundity (more eggs) and lower age of first reproduction (faster sexual maturity) indicate better adult performance.

First, to analyse the effect of higher or lower developmental temperature on adult performance (measured at control conditions, 18°C), two comparisons were performed: 14-18 *vs*. 18-18, for lower developmental temperature; and 26-18 *vs*. 18-18, for higher developmental temperature. Second, to test the cumulative effect on adult performance of exposing flies to both (lower or higher) development and adult temperature, the following comparisons were done: 26-26 *vs*. 26-18, for higher temperatures; and 14-14 *vs*. 14-18, for lower temperatures. If a negative cumulative effect occurs, we expect that individuals exposed to 18°C as adults will have a better performance when compared to those kept at the more extreme temperature, *i.e*., 26-18 (or 14-18) flies will have higher fecundity and lower age of first reproduction than 26-26 (or 14-14) flies. Comparisons with control 18-18 conditions allow to assess the extent to which temperature changes imposed by other thermal treatments are stressful or not, and also if these changes are cumulative.

The normality and homoscedasticity assumptions for analysis of variance were checked and were met in our dataset.

All statistical analyses were performed in R v3.5.3, with the lme4 (Bates et al., 2015), car (Fox and Weisberg, 2019) and lawstat (Hui et al., 2008) packages.

## 3. Results

Thermal plasticity was observed for both age of first reproduction and fecundity – when considering all thermal treatments (Figure 2 and Table A1, significant factor Temp). In general, adult performance was lower for individuals developed at the highest temperature tested (26°C) – Figure 2.

**Figure 2.**
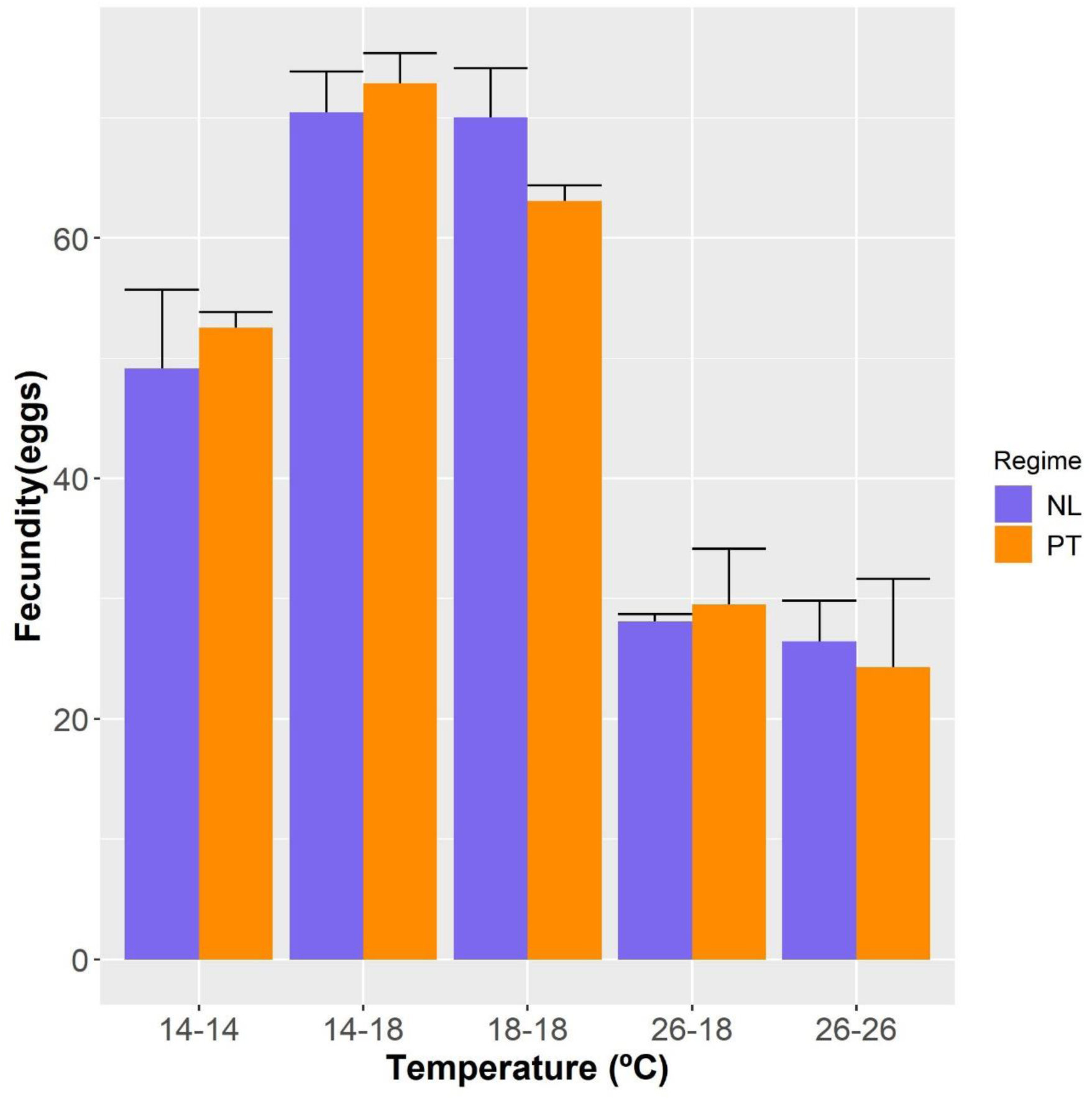
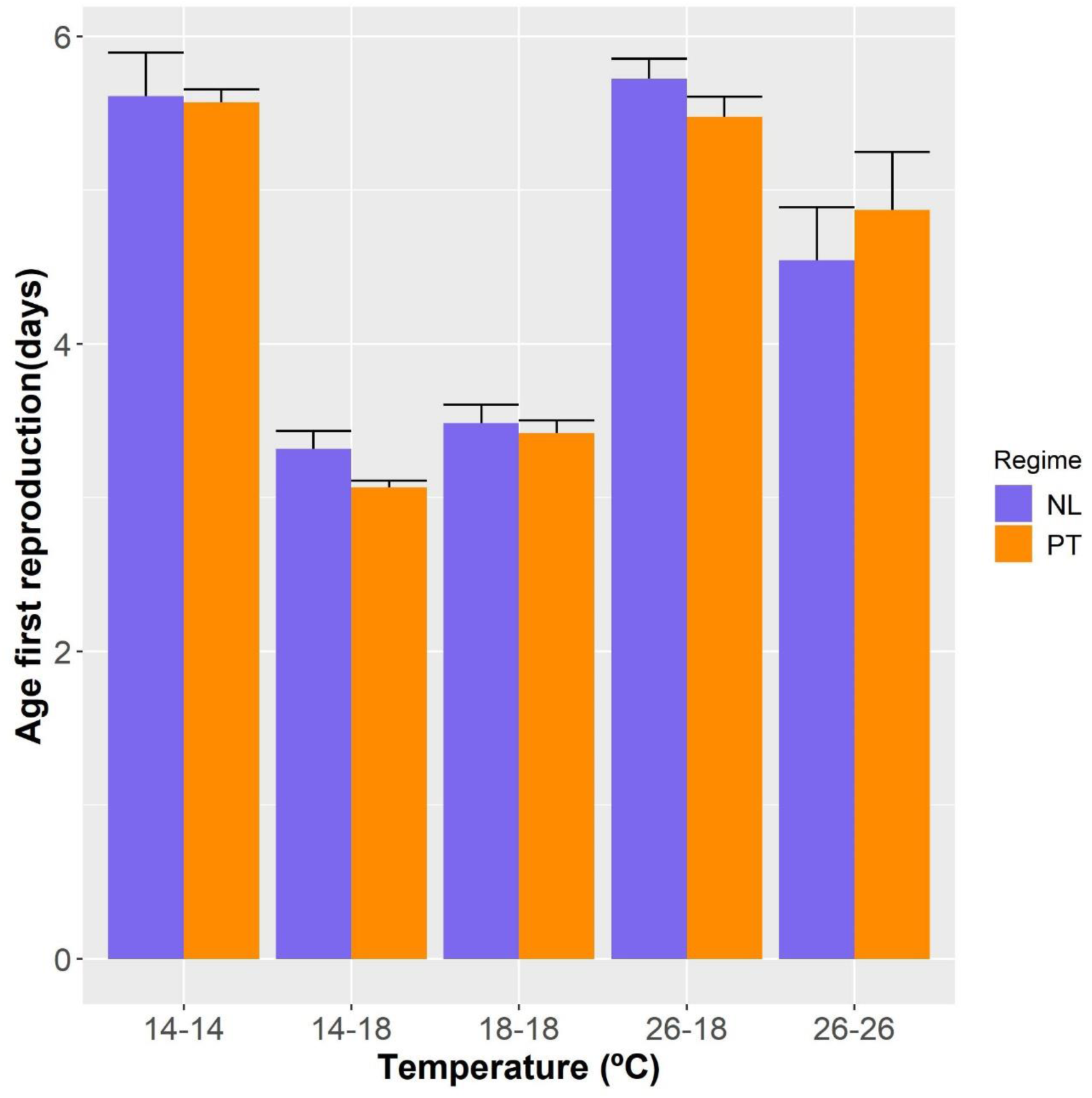
Reproductive performance for individuals exposed to the five different thermal treatments. a) Age of first Reproduction; b) Fecundity (days 7 to 8). Error bars represent the standard error of the 3 replicate populations of each latitudinal population.

The pairwise comparisons to test for the effects of higher or lower developmental temperature on adult performance are described in Table 1. On the one hand, there was a significantly lower adult performance (lower fecundity, higher age of first reproduction) of flies developed at 26°C relative to those developed at 18°C, when tested in the control 18°C environment (see Figure 2, Table 1, 26-18 *vs*. 18-18). On the other hand, individuals developed at 14°C and kept as adults at 18°C reached sexual maturity significantly faster than those always kept at control, 18°C conditions (see Figure 2b, Table 1, 14-18 *vs*. 18-18). No detrimental effect of lower developmental temperature was found for fecundity (see Figure 2a, Table 1, 14-18 *vs*. 18-18).

**Table 1.** Test for the effect of lower (14°C) and higher (26°C) developmental temperature on the thermal response for fecundity characters between populations.

|  | Trait | Model parameters | F<sub>(df1, df2)</sub> |
| --- | --- | --- | --- |
| Higher temperature<br>(26-18 vs. 18-18) | Age of First<br>Reproduction (A1R) | Pop | F<sub>1,4</sub> = 1.792 n.s. |
|  |  | Temp | F<sub>1,4</sub> = 331.78 *** |
|  |  | Pop*Temp | F<sub>1,4</sub> = 0.599 n.s. |
|  | Fecundity | Pop | F<sub>1,4</sub> = 0.769 n.s. |
|  |  | Temp | F<sub>1,4</sub> = 141.89 *** |
|  |  | Pop*Temp | F<sub>1,4</sub> = 1.737 n.s. |
| Lower temperature<br>(14-18 vs. 18-18) | Age of First<br>Reproduction (A1R) | Pop | F<sub>1,4</sub> = 1.641 n.s. |
|  |  | Temp | F<sub>1,4</sub> = 20.891 * |
|  |  | Pop*Temp | F<sub>1,4</sub> = 2.630 n.s. |
|  | Fecundity | Pop | F<sub>1,4</sub> = 0.500 n.s. |
|  |  | Temp | F<sub>1,4</sub> = 3.332 n.s. |
|  |  | Pop*Temp | F<sub>1,4</sub> = 2.791 n.s. |

Note: significance levels: p > 0.05 n.s.; 0.05 > p > 0.01\*; 0.01 > p > 0.001\*\*; p < 0.001 \*\*\*

The test for the cumulative effect of higher developmental temperature showed that switching flies to 18°C (26-18) did not increase performance when compared to keeping them at 26°C in both life stages (26-26); but even decreased in the case of age of first reproduction (see Figure 2 and Table 2, 26-18 *vs*. 26-26), so no negative cumulative effect was found. These results suggest that the negative effect of high developmental temperatures could not be reverted.

**Table 2.** Test for a possible reversion of the effect of lower (14°C) and higher (26°C) developmental temperatures on adult performance

|  | Trait | Model parameters | F<sub>(df1, df2)</sub> |
| --- | --- | --- | --- |
| Higher temperature<br>(26-18 vs 26-26) | Age of First<br>Reproduction (A1R) | Pop | F<sub>1,4</sub> = 0.017 n.s. |
|  |  | Temp | F<sub>1,4</sub> = 15.010 * |
|  |  | Pop*Temp | F<sub>1,4</sub> = 1.574 n.s. |
|  | Fecundity | Pop | F<sub>1,4</sub> = 0.004 n.s. |
|  |  | Temp | F<sub>1,4</sub> = 2.507 n.s. |
|  |  | Pop*Temp | F<sub>1,4</sub> = 0.661 n.s. |
| Lower temperature<br>(14-18 vs 14-14) | Age of First<br>Reproduction (A1R) | Pop | F<sub>1,4</sub> = 0.563 n.s. |
|  |  | Temp | F<sub>1,4</sub> = 398.41 *** |
|  |  | Pop*Temp | F<sub>1,4</sub> = 0.761 n.s. |
|  | Fecundity | Pop | F<sub>1,4</sub> = 0.346 n.s. |
|  |  | Temp | F<sub>1,4</sub> = 61.787 ** |
|  |  | Pop*Temp | F<sub>1,4</sub> = 0.037 n.s. |

Note: significance levels: $p > 0.05$ n.s.; $0.05 > p > 0.01$ \*; $0.01 > p > 0.001$ \*\*; $p < 0.001$ \*\*\*

Conversely, at lower temperatures, performance was reduced only in individuals developed and maintained at such temperatures (see Figure 2, 14-14 *vs*. 18-18). This low performance was reversed when individuals that developed at 14°C were exposed to 18°C as adults: flies from the 14-18 thermal treatment showed significantly higher performance than those from the 14-14 treatment (see Figure 2 and Table 2, 14-18 *vs*. 14-14), showing that the combination of lower juvenile and adult temperatures have a negative effect on adult performance.

No significant differences in thermal plasticity between latitudinal populations were observed, either considering all thermal treatments or in the pairwise tests between them (see Figure 2; Tables 1 and 2, factor Pop*Temp).

## 4. Discussion

We report here that higher – but not lower – developmental temperatures led to lower adult performance in *Drosophila subobscura* flies. These effects were permanent, as they could not be reversed or mitigated by exposing such individuals to a benign environment in the adult stage. In a previous plasticity study with these *D. subobscura* populations, we observed a negative effect of high developmental and adult temperatures on fecundity (Simões et al., 2020). In that experiment, we could not rule out the combined effect of stress, in both life stages, as the tested individuals were kept in their whole life cycle at the same, higher temperature. The present results indicate that there is no combined negative effect of high developmental and adult temperature on reproductive performance. In fact, we observed that flies experiencing higher temperatures across life stages had a higher rate of sexual maturity (younger age of first reproduction) relative to those that only experienced high temperatures during the developmental stage. This is likely due to a faster maturation of females (and, eventually, males) in the imago’s stage due to faster metabolism at higher temperatures (Clarke and Fraser, 2004). This could eventually result in a faster mating speed following emergence, leading to a higher impact on age of first reproduction rather than on fecundity patterns.

We found reduced adult fitness in flies exposed to “non-optimal”, hotter developmental environments, when compared to those developed at control conditions. These “within-generation” carry-over effects have been described in *Drosophila* (Kirk Green et al., 2019; Klepsatel et al., 2019; Porcelli et al., 2017) as well as in other insects (e.g. (Cao et al., 2018; Klockmann et al., 2017; Zhang et al., 2015). Such detrimental effects might result from the irreversible damage of physiological / metabolic pathways and processes, like spermatogenesis and oogenesis, brought upon by stressfully high developmental temperatures. Spermatogenesis has been reported to be particularly vulnerable to heat stress (David et al., 2005). Thus, the longer maturity rate and reduced fecundity observed here might be (at least, partly) due to lower sperm quality / output in males. Also, Porcelli et al. (2017) found that temperatures ∼ 24°C lead to reduced sperm motility in *D. subobscura*. Future analyses should address the extent to which female and male reproductive performances are (differentially) affected by heat stress in these populations.

We also found that fly development at lower temperatures did not reduce adult reproductive performance and, in the case of age of first reproduction, it even led to a better performance relative to individuals raised in control conditions; this may be due to a higher ovariole number in individuals raised at lower temperatures (Moreteau et al., 1997). Yet, this positive effect was not observed when flies were more sexually mature, near their peak fecundity. On the other hand, performance was reduced when individuals were developed and kept as adults at a lower temperature. The fact that fecundity under control conditions (18°C) was similar in flies developed at 18°C or developed at colder conditions, suggests that the lower adult performance at 14°C was a result of a reduction in metabolic rate in adults kept at that temperature. This might reduce oogenesis and lead to lower fecundity even if ovariole number increased.

In a recent study, we found evidence for cold but not heat acclimation in fecundity patterns, with individuals developed at lower temperatures (14°C) having higher fecundity at 14°C than those developed in control conditions – 18°C (Simões et al., 2020). At that point, we could not exclude that such pattern resulted, at least in part, of a general better performance across different environments of flies raised at lower temperatures. The present study indicates that development at lower temperatures does not always lead to improved adult performance, as adult temperature also plays an important role in this case. These results are in contradiction with the “cold is better” hypothesis of developmental plasticity, which posits that individuals developed at colder temperatures have always higher adult performance than individuals raised at other temperatures, i.e. regardless of the test temperature in the adult stage (Huey et al., 1999). With this body of data, we can now rule out that hypothesis, at least in the case of fecundity, as individuals developed at lower temperatures did not show increased fecundity when compared to those developed in control conditions.

Adaptation to different thermal environments is expected to result in differential thermal plasticity between populations (Angilletta, 2009). Porcelli et al. (2017) reported geographical differentiation in the response to heat stress in *Drosophila subobscura*, with northern populations presenting lower viability and fertility. Previously, we found significant differences between the same latitudinal populations studied here, with a higher reproductive performance of the southern (PT) populations when subjected to lower temperatures in both the development and adult stages (Simões et al., 2020). Here, we did not observe any evidence for historical differences in response to lower temperatures. One explanation might be that in Simões et al. (2020), such differences were detected in early fecundity, *i.e*. fecundity tested during the first week of life, while in this study we focused solely on fecundity patterns near peak reproduction, *i.e*. between days 7 and 8. It is possible that the initial differences in fecundity between populations became diluted with time as exposure to colder conditions in adults might potentiate population differences in the rate of sexual maturation, which will reflect more on the initial amount of laid eggs (i.e. early fecundity).

It is an expectation that differences in reproductive performance between thermal treatments might be mediated by body size, since higher developmental temperatures lead to lower body sizes (Kingsolver and Huey, 2008). In a previous plasticity study with these same populations (four generations earlier), we analysed whether variation in wing size accounted for the differences in fecundity across populations and treatments. We concluded that wing size (used as a proxy for body size) was unlikely to be an important factor generating such differences in our latitudinal populations.

In summary, we here demonstrate that increasing the developmental temperature of *D. subobscura* populations ∼8°C above control conditions leads to an irreversible negative effect on reproductive performance, regardless of which adult temperature these organisms are subjected. As observed in other studies, this pinpoints the developmental stage as a critical and most vulnerable stage to climate change and associated heat waves (e.g. see (Kingsolver et al., 2011; Klockmann et al., 2017). The low resilience to increased temperatures during this early stage has likely detrimental consequences to population fitness and persistence. In contrast, we show that this species copes well with colder developmental temperatures, with reduction in performance only occurring when lower temperatures are persistent across life stages.

