## Supplementary material for "High developmental temperature leads to low reproduction despite adult temperature": Table A1

### Supplementary Table

Table A1 - Comparisons of thermal response of fecundity characters between populations and thermal regimes (all five thermal treatments included)

| Trait | Model parameters | $F_{(df1, df2)}$ |
|---|---|---|
| Age of First Reproduction (A1R) | Pop | $F_{1,4}$ = 0.156 n.s. |
| | Temp | $F_{4, 16}$ = 65.702 *** |
| | Pop*Temp | $F_{4, 16}$ = 0.699 n.s. |
| Fecundity | Pop | $F_{1,4}$ = 0.012 n.s. |
| | Temp | $F_{4, 16}$ = 65.890 *** |
| | Pop*Temp | $F_{4, 16}$ = 0.658 n.s. |

Note: significance levels: p> 0.05 n.s.; 0.05>p>0.01*; 0.01>p>0.001**; p<0.001 ***
